# Circadian regulation of muscle growth independent of locomotor activity

**DOI:** 10.1101/778787

**Authors:** Jeffrey J. Kelu, Tapan G. Pipalia, Simon M. Hughes

## Abstract

Muscle tissue shows circadian variation, but whether and how the intracellular circadian clock *per se* regulates muscle growth remains unclear. By measuring muscle growth over 12 h periods, here we show that muscle grows more during the day than at night. Inhibition of muscle contraction reduces growth to a similar extent in day and night, but does not ablate the circadian variation in growth. Muscle protein synthesis is higher during the day compared to night, whereas markers of protein degradation are higher at night. Mechanistically, the TORC1 inhibitor rapamycin inhibits the extra daytime growth, but no effect on muscle growth at night was detected. Conversely, the proteasomal inhibitor MG132 increases muscle growth at night, but has no effect during the day, irrespective of activity. Ablation of contractile activity rapidly reduces muscle protein synthesis both during the day and at night and leads to a gradual increase in Murf gene expression without ablating circadian variation in growth. Removal of circadian input by exposure to either permanent light or permanent darkness reduces muscle growth. We conclude that circadian variation in muscle growth is independent of the presence of, or changes in, physical activity and affects both protein synthesis and degradation in distinct circadian phases.

## Introduction

The need to adapt to the daily cycle of light and dark drove evolution of the circadian intracellular molecular clock (Gerhart-Hines and Lazar, 2015; Paranjpe and Sharma, 2005; Vaze and Sharma, 2013). The animal circadian clock mechanism is highly conserved, but its output has adapted dramatically as species occupied their niches (Paranjpe and Sharma, 2005). An obvious animal circadian rhythm is day/night difference in locomotor activity, which is powered by contraction of skeletal muscle. Roles for the circadian biological clock as a contributing factor of muscle growth and/or performance have been revealed (Chtourou and Souissi, 2012; Grgic et al., 2019; Teo et al., 2011). However, a long-standing question, particularly in physiotherapy and sports medicine, is whether time-of-day affects muscle growth either intrinsically, or in response to exercise. Studies seeking the best time-of-day to exercise to build muscle remain largely inconclusive (Hatfield et al., 2016; Küüsmaa et al., 2016; Sedliak et al., 2009, 2008, 2007; Souissi et al., 2002). In such studies it has been difficult to dissociate the intrinsic clock from other circadian oscillations, such as locomotor activity or food intake. One confounder is that human subjects can exert higher muscle force in the evening than in the morning (Chtourou and Souissi, 2012; Grgic et al., 2019; Teo et al., 2011). Thus, the question of what role the circadian clock *per se* has in the regulation of muscle mass and physiology remains open. Our aim here is to test the hypothesis that the intrinsic circadian clock in muscle regulates muscle growth independent of locomotor activity.

Circadian rhythms are the result of an endogenous biological clock that oscillates with a >24 h-period (Czeisler et al., 1999) interacting with environmental cues or cycles resulting from Earth’s rotation that entrain the clock to a 24 h-period. It is well-known that muscle growth promoting factors such as hormonal, feeding and locomotor activities are under direct control by the circadian clock (Huang et al., 2011). Recent studies in human suggest that disturbance of the circadian rhythm, via shift work, lack of sleep and social/travel jetlag, affect skeletal muscle homeostasis (Aisbett et al., 2017; Lucassen et al., 2017; Roenneberg et al., 2012; Yu et al., 2015). Around 70% of the European population from industrialised cities was found to suffer from ‘social jetlag’ i.e. discordance between the circadian clock (i.e. sleep-wake cycle) and social activities (Roenneberg et al., 2012). Social jetlag may lead to chronic sleep loss and contribute to the development of weight-related pathologies i.e. obesity (Roenneberg et al., 2012). Altered muscle metabolism can contribute to obesity (Collins et al., 2018; Wells et al., 2008). In addition, sleep deprivation or restriction impairs acute maximal muscle strength in compound movements (Aisbett et al., 2017), which could diminish muscle maintenance. Poor sleep quality and later sleeping timing are risks factors for the development of ageing-related muscle wasting (sarcopenia) in middle-aged individuals (Lucassen et al., 2017). Furthermore, a higher prevalence of diabetes, metabolic syndrome, and sarcopenia was found in the evening *versus* morning chronotype among middle-aged individuals (Yu et al., 2015). These studies, therefore, suggest a potential link between the circadian clock machinery, skeletal muscle physiology and population health. Nonetheless, a direct relationship between the dysregulation of circadian clock and loss of muscle and strength has yet to be established.

It is not only adults that may be affected. Altered circadian rhythm during pregnancy or in the young could, if it affected muscle building, have lifelong effects because correlative data increasingly associates low birth weight (a major component of which is reduced muscle mass) with a range of adult medical problems (Belbasis et al., 2016; Jornayvaz et al., 2016). One major plausible causal link is an alteration in muscle/fat balance in early life.

Studies in mice have demonstrated that skeletal muscle itself possesses a robust circadian rhythm with >1500 genes that oscillate expression over a 24 h period (McCarthy et al., 2007; Miller et al., 2007; Pizarro et al., 2013; Zhang et al., 2014). Many of these genes are involved in metabolism, transcription and signalling in muscle (Hodge et al., 2015; McCarthy et al., 2007). Whole-body knockout of the core clock gene *bmal* results in progressive and dramatic muscle atrophy, decreased total activity level, decreased muscle force, myofibrillar disorganization, and premature aging (Andrews et al., 2010; Kondratov et al., 2006), but recent inducible muscle-specific *bmal* knockout failed to recapitulate these phenotypes (Dyar et al., 2016, 2014; Yang et al., 2016), suggesting that muscle defects in the whole-body *bmal* knockout were secondary to clock disruption in other tissues. Two circadian rhythms likely to mediate such secondary effects by other tissues are 1) neurally-driven muscle activity, and 2) food intake, which alters gut/liver-derived nutrition. We recently developed a method of measuring muscle growth over a period of a few hours in the live zebrafish (Hinits et al., 2011; Roy et al., 2017). Here we use this zebrafish model to separate the roles of the circadian clock and physical activity in muscle anabolism and catabolism leading to growth.

Zebrafish (*Danio rerio*) offer advantages for the study of muscle circadian clock (Gibbs et al., 2013). First, like people, but unlike rodents, zebrafish are diurnal, being active in day but less active at night (Cahill et al., 1998; Hurd et al., 1998; Hurd and Cahill, 2002). Most muscle clock studies to date have been conducted using laboratory mice (*Mus musculus*). Mice are nocturnal, being active mainly at night and have fragmentated daytime sleeping, contrasting with human and zebrafish. Analysis of circadian gene expression profiles in baboons (*Papio anubis*), a primate relative of humans, and mice indicated that, as expected, the peak phase of expression of the core clock components (e.g. *bmal* and *per1*) was shifted by ∼12 h between baboon and mouse tissues (Mure et al., 2018). Nevertheless, a consistent ∼12 h difference in the peak phases of expression was not observed among the common cycling mRNAs between baboon and mouse in the different tissues. For example, in heart 47% of the shared rhythmic genes cycled with <6 h phase difference, whereas in cerebellum 65% of shared cycling genes showed a 10-15 h phase difference (Mure et al., 2018). This shows that the diurnal-nocturnal changes downstream of the core clock components are complex and species- or tissue-specific. There is thus a need to look beyond rodents when modelling the muscle clock. Second, like all animals, zebrafish larvae show circadian changes in motor activity (Cahill et al., 1998; Hurd and Cahill, 2002). While it has long been known that exercise/activity can induce muscle growth, it has not been clear whether circadian differences in muscle growth rate result from differences in activity pattern or due to the clock *per se*, e.g. the circadian expression of clock-controlled genes. To circumvent the effects by activity, muscle growth can be studied in zebrafish after anaesthesia or genetic mutation of the acetylcholine receptor (*fixe* mutant), which render larvae immotile and this affects myogenesis (Yogev et al., 2013). Third, the zebrafish larva does not feed until 5 days post-fertilisation (5 dpf) (Strähle et al., 2012), so the other prominent confounding effect of feeding, a strong circadian entrainment cue (zeitgeber) (Davidson and Stephan, 1999; Shavlakadze et al., 2013), can be eliminated. Fourth, zebrafish are transparent and all zebrafish cells (including muscle cells) are directly entrainable by light, meaning that the circadian clock can operate cell-autonomously (i.e. intrinsically) in individual cells, and that a central oscillator (e.g. the superchiasmatic nucleus (SCN) or pineal gland) is not required for light entrainment (Whitmore et al., 2000). In people, the amplitude of fluctuations in circadian-controlled mRNAs was reduced and gradually lost in biopsy muscle cell cultures, perhaps due to lack of entrainment from SCN rhythm (Hughes et al., 2009; Perrin et al., 2018). Thus, compared to mammalian *in vivo* or muscle cell culture studies, zebrafish larvae enable study of the role of the circadian clock during muscle growth, independent of exercise and feeding, which has not been possible in other model systems.

Here we set out to examine the role of circadian rhythm in skeletal muscle growth in the embryonic and larval zebrafish. We show that muscle grows best in a circadian 12 h-light/12 h-dark (LD) regime, and that muscle grows more during day than night. In the absence of physical activity muscle growth continues to show circadian variation. Physical activity augments muscle growth in both dark and light phases. Analysis of anabolic and catabolic pathways shows that anabolism is greatest in daytime, is promoted by physical activity and correlates with TORC1 activity. Rapamycin, a TORC1 inhibitor, specifically reduced muscle growth in the day. In contrast, catabolism is greatest at night, correlates with accumulation of Murf mRNAs and MG132, a proteasome inhibitor, specifically enhanced growth at night. Catabolism is promoted by the intrinsic circadian clock and is partially overcome by physical activity. We conclude that muscle metabolism and growth are regulated by the circadian clock and the circadian difference is independent of the effect of physical activity.

## Results

### Consistent myotome growth despite individual variation in size

We have developed a robust volumetric muscle growth assay that can detect growth in real time in the living zebrafish larva, selecting somite 17 because of its ease of imaging at the trunk-tail boundary (Hinits et al., 2011; Roy et al., 2017) (Fig. S1). The volume of the myotome of somite 17 grows by around 170% between 1-3 days post-fertilisation (dpf), and 40% between 3-5 dpf, (Fig. 1A). Fibre numbers increase in parallel, but the major component of growth is increase in fibre size (Fig. S2A,B). Individuals vary in myotome volume both within and between lays (Figs 1B and S2C). When single unmanipulated fish were measured repeatedly at 3 dpf and 4 dpf, all grew by a similar amount, although final size correlated with initial size (Fig. S2C). We conclude that myotome volume can be accurately measured, that comparisons between individuals are best made within lays and that growth can be most accurately assessed by repeatedly measuring single individuals.

**Figure 1.**
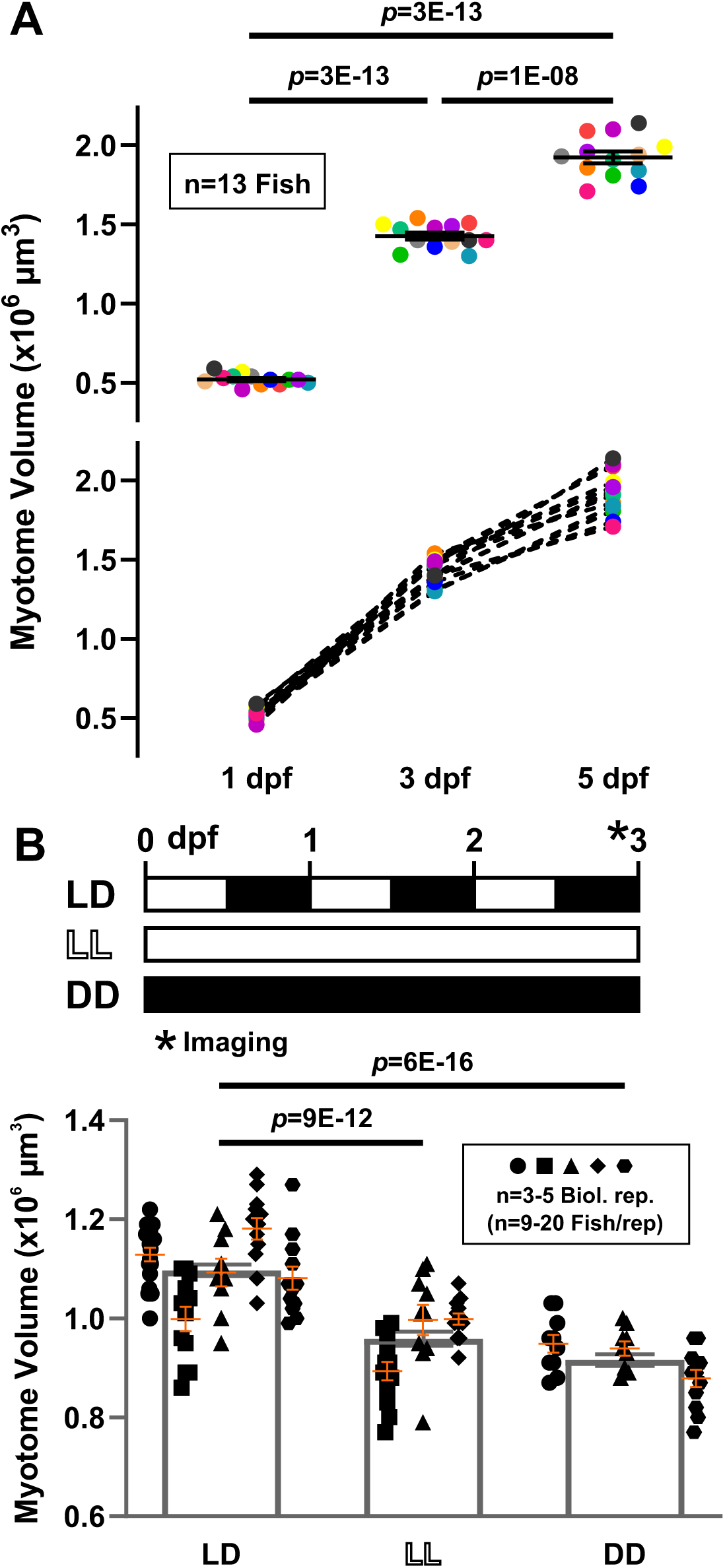
Effect of altered light regimes on the volumetric growth of zebrafish larval myotome. **A.** Two representations of growth of myotome in somite 17 between 1 and 5 dpf. Colours represent different individual fish from a single lay (n=13 fish). **B.** Sibling larvae collected within 1-2 h of fertilization were reared under different light regimes, LD, LL, or DD as schematised above. Myotome volume was measured at 3 dpf (n=29-63 fish, from 3-5 biological replicates). Symbols represent distinct lays (biological replicates). Orange bars represent mean ±SEM, black boxes represent global mean. Note lay-to-lay variation.

### Light/dark circadian cycle promotes muscle growth

We first examined the effects of circadian illumination on muscle growth in post-hatch larvae aged 3 dpf, a stage before independent feeding, when the larvae would naturally be living in hiding and using their muscle to escape predators or noxious environmental factors. In a 12 h light: 12 h dark (LD) regime, lays varied in mean myotome volume between 1 and 1.2 × 10^6^ μm^3^ (Fig. 1B). When siblings were exposed to either constant light (LL) or constant darkness (DD) from just after laying their myotome volume was reduced by around 15% (Fig. 1B). Thus, a normal circadian light cycle is required for optimal myotome growth.

### Muscle grows more in day than night

To assess circadian effects in more detail, the growth of the myotome in 3 dpf LD fish was measured every 12 h over a 24 h period. The myotome grew about 50% faster during the day than at night, whether measured in absolute or percentage terms (Figs 2A and S3A), and the growth difference was consistent across different lays and experiments (Fig. S3B). This day/night variation correlates with the rhythmic expression of the molecular clock; expression of *bmal1a* (one of the core clock genes) fluctuates in a rhythmic manner between 3-5 dpf, peaking late in the day (Fig. S3C)(Dekens and Whitmore, 2008). Measurement of growth every 12 h over a 48 h period between 3-5 dpf consistently shows a growth difference between day and night, although overall growth is slower on day 4 than on day 3 (Fig. S3D). To distinguish whether the slower growth at night simply reflected the slowing of growth with developmental stage, the light cycle was reversed on sibling larvae (DL) and their myotome growth measured (Fig. 2B). The larval myotome consistently grew more in the 3.5-4 dpf light phase than in the 3-3.5 dpf dark phase. We conclude that the larval zebrafish myotome grows more in the day than in the night, irrespective of developmental stage.

**Figure 2.**
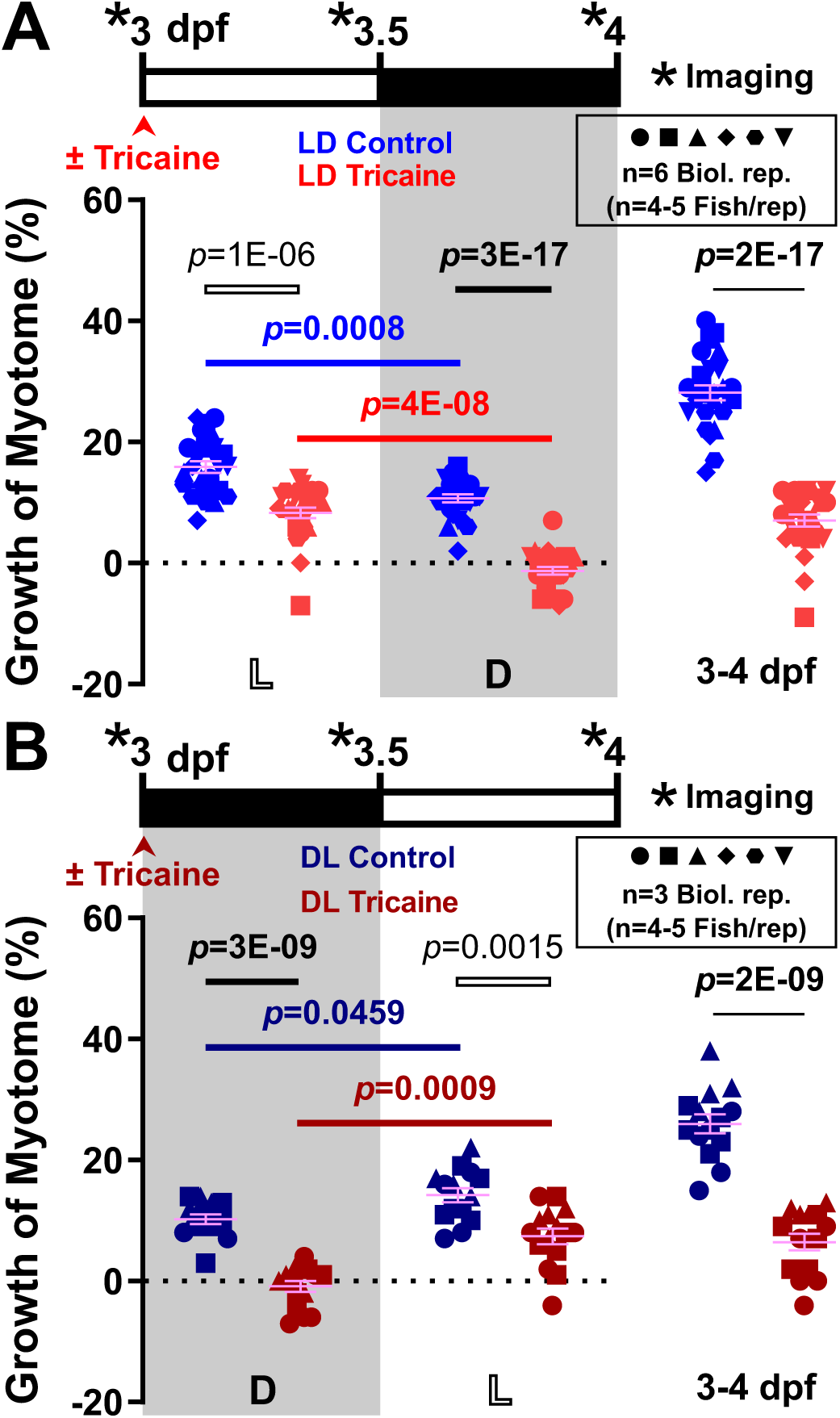
Circadian differences in volumetric growth of the myotome and motor activity of larval zebrafish. Larvae were raised under LD (**A**; n=25-26 fish from 6 biological replicates) or DL (**B**; n=14-15 fish from 3 biological replicates) and were either untreated (Control, blue) or anaesthetised (Tricaine, red) from 3-4 dpf. Confocal imaging was performed every 12 h (*) over the 24 h period to measure myotome volume. Graphs show growth of myotome (%) of individual fish over each 12 h period (left and centre) or the full 24 h (right). Complete data on the same LD fish are shown in Fig. S3A,B. Symbols shapes distinguish biological replicates from separate lays. Colours represent different treatments. L light, D dark.

### Circadian differences in growth independent of activity

A possible explanation for the circadian difference in muscle growth is differing levels of physical activity between day and night. Larvae spontaneously swim more during the day, raising the possibility that activity promotes muscle growth (Cahill et al., 1998; Hurd and Cahill, 2002). We therefore grew larvae in the anaesthetic tricaine from 3-4 dpf and measured myotome growth (Figs. 2A,B and S2A,B). We have previously shown that tricaine blocks movement by inhibiting neural activity (Attili and Hughes, 2014). Growth was significantly reduced by inactivity, both during the day and night. A clear circadian difference remained, however, between growth during the day and night (Figs. 2A,B and S2A,B). The myotome of inactive fish grew during the day but failed to grow, and even showed a tendency to shrink, at night (Fig. 2A). Importantly, such a day/night growth difference was shown to remain consistent over a 48 h period in anaesthetised fish (Fig. S3D). When the light cycle was reversed, again, the myotome of anaesthetised fish grew in the later light phase and shrank in the earlier dark phase (Fig. 2B). These data show that circadian effects on myotome growth are independent of muscle electrical and contractile activity. Interestingly, the reduction in growth caused by inactivity was quantitatively similar between day and night (Fig. 2B), suggesting that the amount of activity-dependent growth is not proportional to the quantity of activity *per se*, which differed substantially between the light and dark phases (Cahill et al., 1998; Hurd and Cahill, 2002).

### Both circadian clock and physical activity promote anabolism

To determine how activity regulates growth, we next analysed protein synthesis in muscle by using O-propargyl-puromycin (OPP) (Horisawa, 2014) to compare the quantity of ribosomes translating mRNA in different conditions. OPP incorporation was performed at different zeitgeber time (ZT). During the early day (ZT0-2), OPP labels muscle well above the control background level revealed by cycloheximide-treatment before OPP (Fig. 3A,B). Upon tricaine treatment, labelling dropped significantly (Fig. 3A,B). OPP incorporation remained above the cycloheximide control in the presence of tricaine however (Fig 3A-C). We conclude that activity promotes muscle protein synthesis, but that some protein synthesis continues during the day in the absence of all muscle activity.

**Figure 3.**
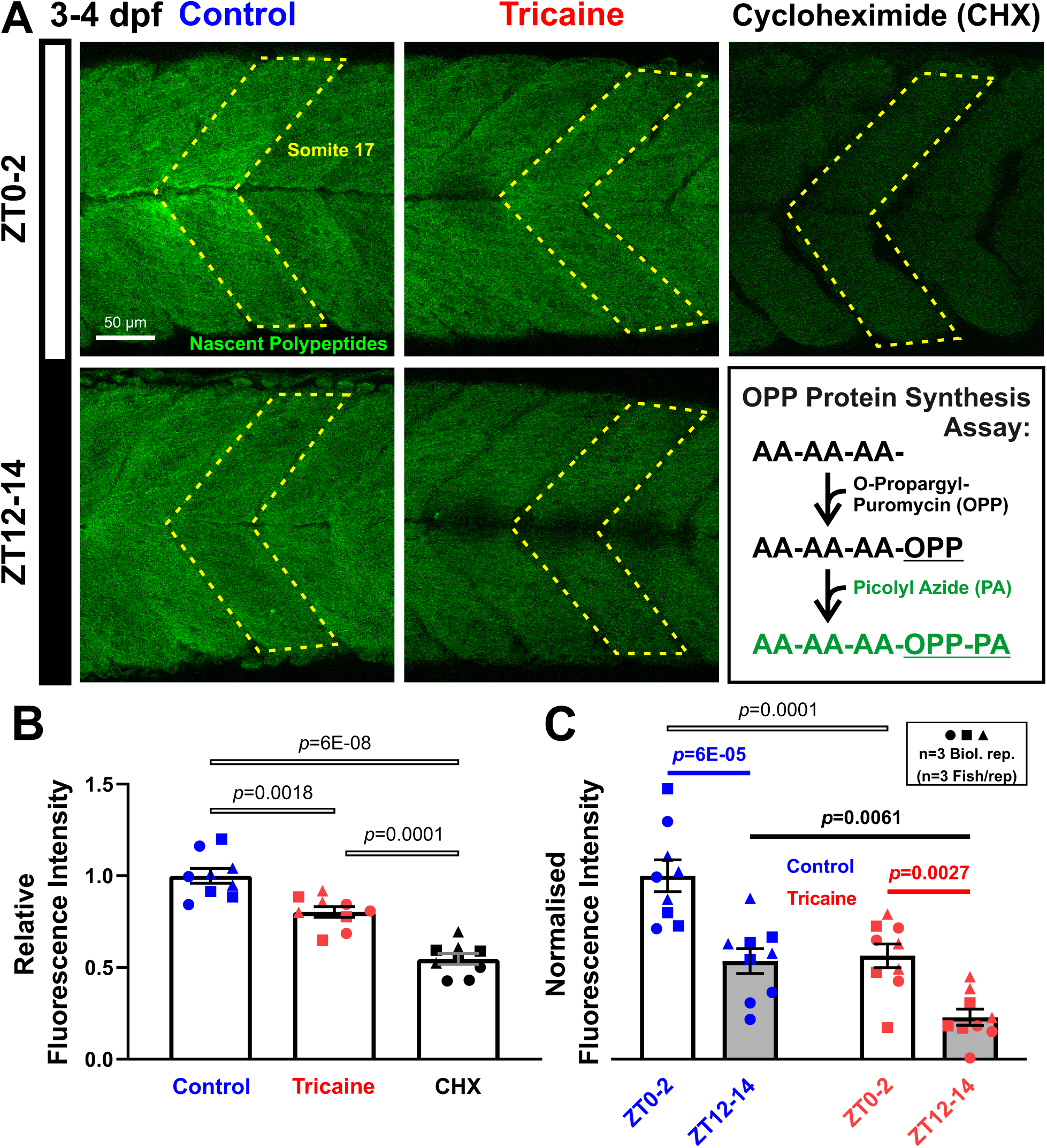
Circadian differences in nascent protein synthesis in larval zebrafish. **A.** Nascent proteins visualized by confocal microscopy in larvae that were raised under LD either untreated (Control) or anaesthetised (Tricaine) at 3 dpf, and then treated with O-propargyl-puromycin (OPP) for 2 h from ZT0-2 or ZT12-14. Inset schematizes amino acid (AA) chain termination by OPP and detection with picolyl azide (PA). Larvae treated with cycloheximide (CHX) and OPP from 3 dpf were analysed at ZT2 as a negative control. **B**,**C.** Quantification of OPP-PA fluorescence of somite 17 (n=9 fish from 3 biological replicates). Signals were normalised to Control ZT0-2 samples (B), and after subtraction of CHX background (C). Colours represent different treatments. Symbols represent distinct lays (biological replicates).

We next asked whether protein synthesis showed circadian variation in active fish. Strikingly, OPP incorporation was reduced in active fish at night (ZT12-14; Fig. 3A,C). The extent of reduction in OPP incorporation between morning and evening was comparable to the effect of tricaine in the morning. To determine whether reduced activity at night caused the reduction in protein synthesis, OPP incorporation was assessed in tricaine-treated larvae at ZT12-14. Tricaine reduced OPP incorporation still further, such that a difference in OPP incorporation persisted between day and night in the complete absence of activity (Fig. 3A,C). Thus, the circadian clock promotes muscle protein synthesis preferentially during the light phase, and physical activity promotes muscle protein synthesis both during day and night, paralleling their effects on volumetric growth.

### Activity and the circadian clock act upon TORC1

Muscle protein synthesis is regulated by TORC1 signalling in response to various hypertrophic stimuli (Yoon, 2017). To determine if developmental muscle growth was regulated similarly, we first examined a sensitive read-out of TORC1 activity in zebrafish larvae (Yogev et al., 2013), phosphorylation of ribosomal protein S6 (Figs 4A and S4). pS6/S6 ratio showed circadian variation, being high during the day (ZT3 and ZT9) and decreasing at night (ZT15 and ZT21) (Fig. 4B). The increased pS6 in early morning was present despite the total S6 protein content being higher at that time. As we have previously shown that a significant fraction of pS6 at earlier stages is in muscle (Yogev et al., 2013), the data suggest that TORC1 activity is elevated in muscle during the day compared to the night.

**Figure 4.**
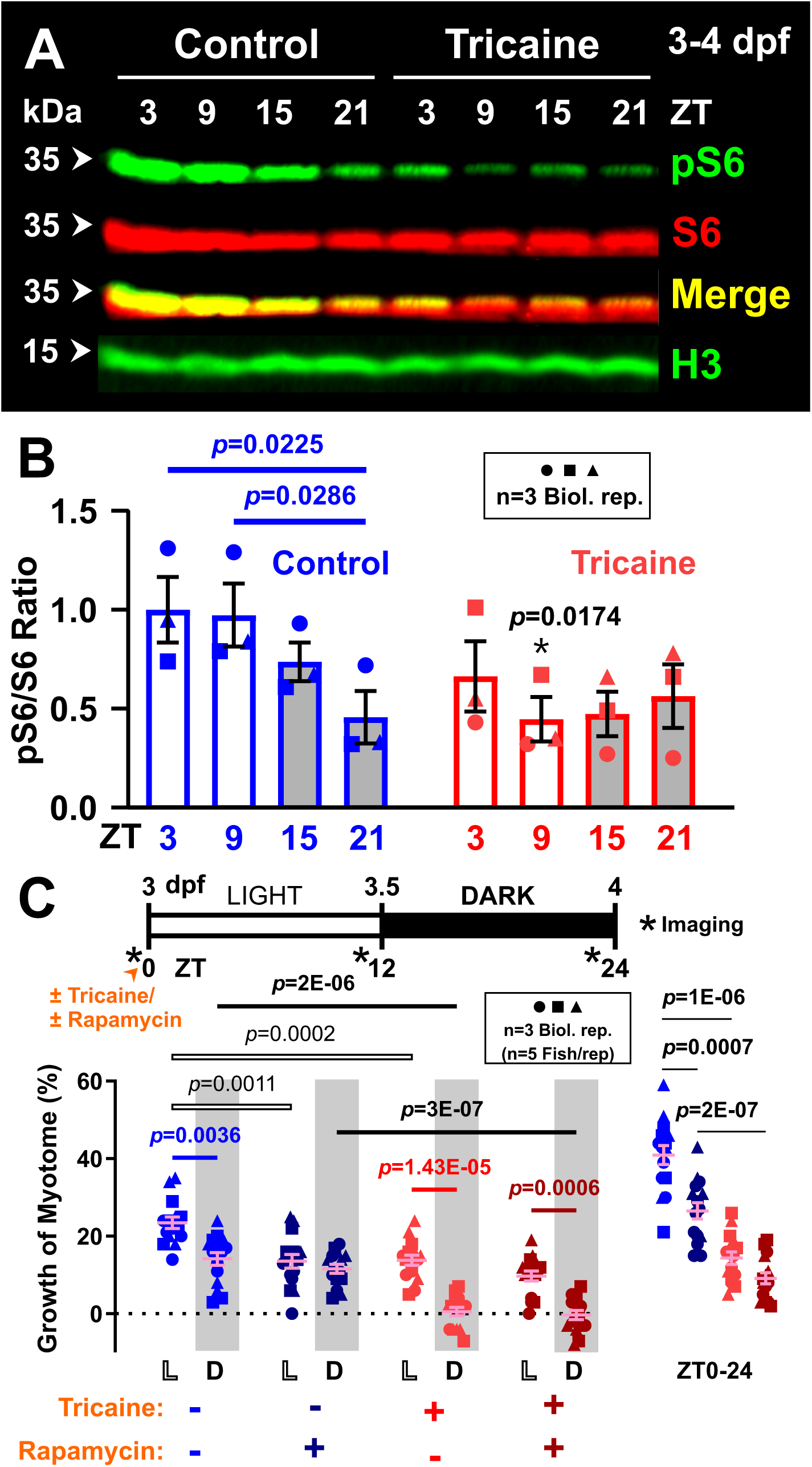
Circadian differences in the activity of mTOR signalling pathway in larval zebrafish. **A.** Western analysis of phosphor-Ribosomal protein S6 (pS6), total RpS6 (S6) and Histone H3 loading control (H3) in larvae raised under LD that were either untreated (Control) or anaesthetised (Tricaine) from 3-4 dpf. Total protein was extracted from whole larvae every 6 h over a 24 h period, at ZT3, ZT9, ZT15, and ZT21. **B.** Quantification of pS6/S6 ratio (n=3 biological replicates, shown in Fig. S4). Ratios were normalised to control samples at ZT3. **C.** Growth of the myotome (%) in larvae raised under LD that were either unanaesthetised (Control, blue) or anaesthetised (Tricaine, red) from 3-4 dpf. The larvae were also treated with either DMSO vehicle (light colours) or 10 μM rapamycin (deep colours). Symbols represent distinct lays (n=15 fish, from 3 biological replicates). L light; D dark.

We next tested the hypothesis that physical activity regulates TORC1 activity. Tricaine treatment began to reduce pS6/S6 ratio after 3 h of treatment, but had a large inhibitory effect after 9 h at ZT9 (Fig. 4A,B). In contrast, tricaine had no significant effect on pS6/S6 ratio during the dark phase. This result is markedly different from the reduction in volumetric muscle growth caused by tricaine during the dark phase (Fig. 2A). Nevertheless, tricaine did appear to affect ribosome biogenesis, as the raised level of total S6 protein present in the morning was reduced by tricaine treatment (Fig. 4A). These experiments suggest that TORC1 activity is higher during the day than night and that the diurnal rise in TORC1 activity depends on muscle contraction.

### Circadian clock-dependent role of TORC1 in myotome growth

To examine the role of TORC1 in myotome growth in more detail, we measured muscle growth during the light and dark phases after inhibiting TORC1 with rapamcyin, which we have previously shown to rapidly reduced pS6/S6 ratio in larval zebrafish (Yogev et al., 2013). As before, vehicle-treated control fish grew more during the day than during the night (Fig. 4C blue symbols). Treatment with rapamycin reduced growth only during the day, but had no effect on growth at night (Fig. 4C deep blue symbols). As shown previously, tricaine treatment reduced growth in both phases (Fig. 4C, red symbols). Adding rapamycin to tricaine-treated fish appeared to reduce growth slightly during the day, albeit not reaching statistical significance. However, rapamycin had no effect at night in anaesthetised fish (Fig. 4C, deep red symbols). Importantly, however, the presence of rapamycin did not prevent the growth induced by activity at night. (Fig. 4C). We conclude that rapamycin-inhibitable TORC1 activity is not required for physical activity to promote muscle growth, at least at night. During the day, however, rapamycin appears to inhibit the ability of physical activity to promote growth.

### Circadian induction of catabolism at night is suppressed by activity

Our finding that physical activity drives anabolism by different mechanisms in day and night raises the question of whether catabolism is also regulated by the circadian clock. Several genes are known to regulate catabolism in muscle tissue by activating proteasomal proteolysis of sarcomeric proteins and this is thought to regulate muscle size (Bodine et al., 2001; Witt et al., 2008). Among such genes are those encoding the Muscle RING Finger (Murf) family of Tripartite Motif (Trim) proteins that have been reported to undergo circadian changes in mRNA level in muscle (Amaral and Johnston, 2012; Dyar et al., 2018, 2015; McCarthy et al., 2007). Analysis of *murf1*/*trim63a* and *murf2/trim55b* mRNA levels in 3-5 dpf larvae by qRT-PCR suggested that these genes undergo circadian cycling, such that they peak in every dark phase at ZT15 (Fig. 5A). The algorithm JTK_Cycle (Hughes et al., 2010) confirmed *murf1*/*trim55b* as a significant cycling component. Moreover, inactivity was observed to enhance the overall level of mRNA for these Murfs. Given that Murfs are known to encode E3 ubiquitin ligases that are linked to muscle protein degradation (Bodine et al., 2001; Witt et al., 2008), these observations raise the possibility that protein catabolism is enhanced in myotomal muscle at night.

**Figure 5.**
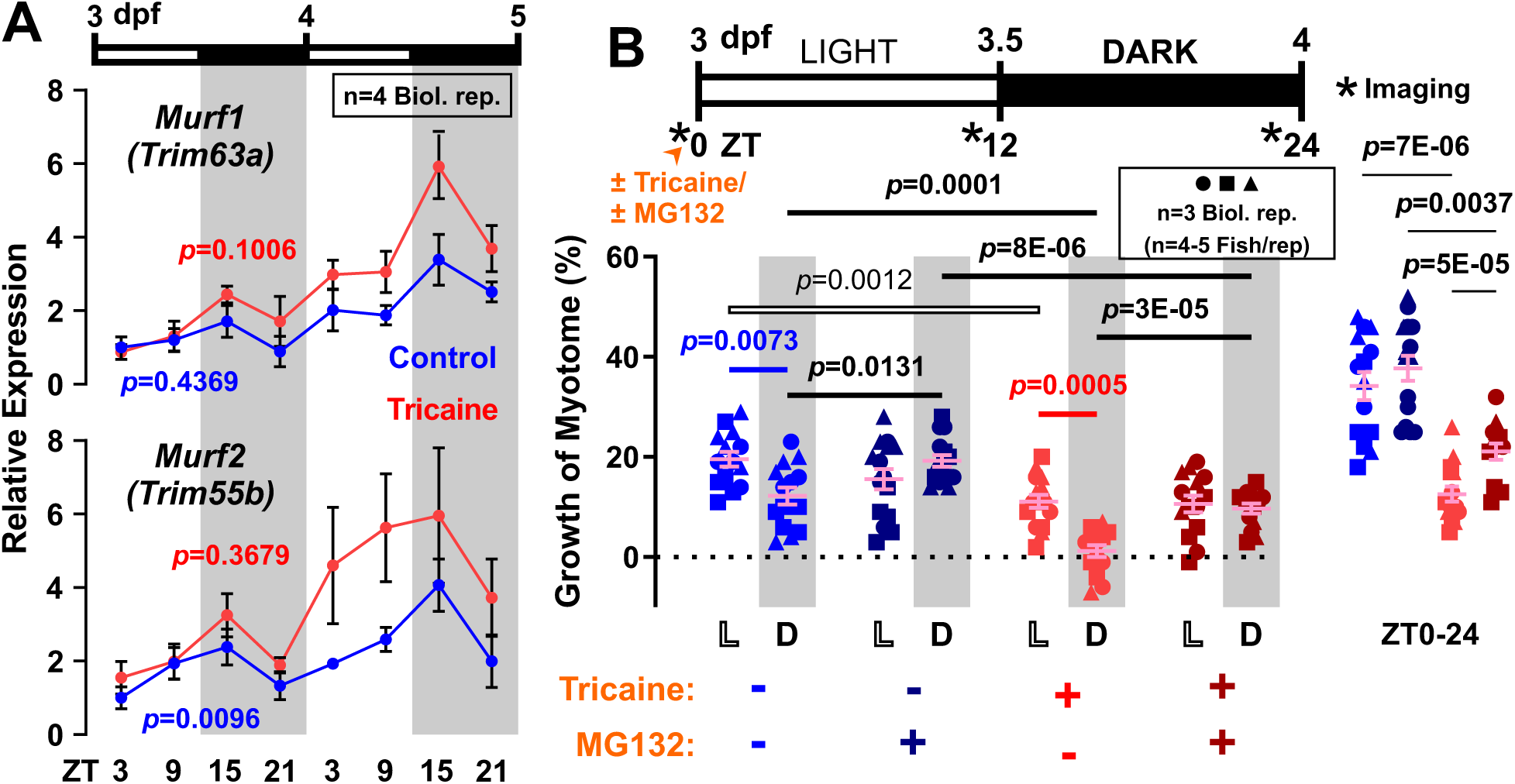
Circadian differences in the expression of atrophic genes in larval zebrafish. Effects of proteasome inhibitor MG132 on the volumetric growth of zebrafish larvae. **A.** Expression of atrophy-related genes *trim63a* and *trim55b*, encoding Murf1 and Murf2, was assayed by qPCR on total RNA collected every 6 h from untreated (Control, blue) and 3-5 dpf anaesthetised (Tricaine, red) larvae raised under LD (n=4 biological replicates). *Ef1a* was used for normalisation. Statistic represents JTK_Cycle (see Methods). **B.** Growth of myotome (%) in larvae raised under LD that were either unanaesthetised (Control, blue) or anaesthetised from 3-4 dpf (Tricaine, red). The larvae were also treated from 3 dpf with either DMSO vehicle (light colours) or 10 μM MG132 (deep colours). Symbols represent distinct lays (n=14-15 fish from 3 biological replicates). L light; D dark.

To address the functional role of protein catabolism in volumetric muscle growth, we next used the proteasomal inhibitor MG132, which we have previously shown to rapidly up-regulate known targets of the proteasome, p53 and Sqstm1, in larval zebrafish (Yogev et al., 2013), to block protein degradation and analysed its effect on circadian muscle growth. MG132 alone had no effect on myotome growth during the day, but increased growth at night (Fig. 5B deep blue compared to blue symbols). As described previously, tricaine reduced growth in both phases (Fig. 5B, red symbols). Strikingly, treatment of inactive fish with MG132 still increased growth at night, but had no detectable effect during the day (Fig. 5B deep red compared to red symbols). We conclude that the circadian clock activates catabolism of muscle proteins at night that counteracts on-going basal growth.

## Discussion

It has previously not been possible to associate circadian differences in muscle protein metabolism with muscle growth over a single 24 h period. Data presented in the current work provides a unified analysis leading to three major conclusions regarding circadian control of muscle growth. Firstly, we show that larval zebrafish muscle grows faster during the day than at night. Secondly, we demonstrate that both protein synthesis and degradation are differentially active in day and night and that manipulation of each process changes muscle growth in a specific circadian phase. Thirdly, we find that the circadian growth difference is independent of both physical activity and food intake. To conclude, we present a testable hypothesis of how the circadian clock interacts with physical activity to control muscle growth.

### Muscle grows more during day than night

Using our zebrafish muscle growth model, volumetric growth can be measured on short timescales in parallel with protein turnover. By performing imaging across day and night, we report a circadian difference in muscle growth in live animals *in vivo* (Fig. 2). To our knowledge, this is the first such report. Growth in the day is approximately double that at night. The higher growth in the day parallels a higher rate of protein synthesis and TORC1 activity (Figs 3,4). Functionally, rapamycin treatment decreased muscle growth in the day (but not at night), and this removed the growth difference between day and night (Fig. 4). This suggests that maximal growth in the light phase requires TORC1 activity. On the contrary, MG132 specifically augmented growth at night (but not in the day), suggesting that proteasome activity in the dark phase limits muscle growth (Fig. 5). The lower level of protein synthesis and TORC1 activity (Figs 3,4) and higher *murf1* and *murf2* expression at night (Fig. 5), indicate that the slower growth in the dark phase arises from a combination of decreased anabolism and increased catabolism.

In adult muscle, mass maintenance is also thought to be controlled by protein turnover, which is suggested to be regulated by the circadian clock. Previous studies, generally performed on adult rodent limb muscle, indicated that protein synthesis, as reflected by either p70S6K and ERK1/2 phosphorylation (Chang et al., 2016) or puromycin incorporation (Dyar et al., 2018), display a day/night variation in skeletal muscle. In addition, the expression of atrogenes, such as *atrogin1* and *murf1*, were shown to undergo rhythmic circadian changes in skeletal muscle (Dyar et al., 2018, 2015; McCarthy et al., 2007; Perrin et al., 2018). Some of these circadian differences were shown to be abolished by loss-of-function of core clock genes (Dyar et al., 2018, 2015), indicating the critical requirement of the molecular clock in regulating muscle metabolism. However, a functional link between these cycling metabolic activities and circadian change in muscle mass remained to be discovered. Our data show that, at least in the myotomal muscle of developing zebrafish, rapamycin-sensitive TORC1 enhances muscle growth in the day, whereas proteasomal activity suppresses muscle growth at night.

### Circadian regulation of muscle growth is independent of physical activity

No previous study has been able to examine muscle growth in the absence of circadian variation in muscle contractile activity. We show that faster growth in day than at night is a circadian property that is independent of locomotor activity (Fig. 2). Inhibition of all muscle activity decreased growth in both day and night phases similarly, strengthening the view that physical activity enhances growth in developing zebrafish (Yogev et al., 2013). Nevertheless, inactive muscle maintains a circadian growth difference that is manifested by net growth in the day but no growth at night (Fig. 2). Thus, the difference in growth between day and night is just as great in inactive muscle as in active muscle. This argues that the circadian clock directly controls muscle growth independent of physical activity.

Zebrafish larvae normally exhibit a diurnal lifestyle (Cahill et al., 1998; Hurd and Cahill, 2002). Nevertheless, swimming enhances muscle growth equally in day and night phases. It therefore appears that even the low levels of activity observed at night are sufficient to maintain a good rate of activity-dependent growth.

In many circadian studies it is difficult to eliminate rhythmic feeding as a potential source of circadian variation in metabolism (Davidson and Stephan, 1999; Shavlakadze et al., 2013). We eliminate such feeding cues in several ways. Firstly, larval zebrafish do not begin to feed until 5 dpf as their jaws are not sufficiently developed. Secondly, no food was provided for larvae in our study. Thirdly, abolition of motor activity prevented any feeding behaviours. At the stages examined, presumably nutrition and substrates for muscle anabolism derive from the yolk via the circulation (Fraher et al., 2016); whether nutrient availability shows circadian variation in larvae is unknown. Thus, our data do not as yet address the question of whether the muscle-intrinsic clock or extrinsic signals from a central clock drive circadian variation in muscle growth.

Our study has chiefly focused on the fourth day of development because this stage is after hatching but before independent feeding. It is also a stage at which general anaesthetic can be administered for 24 h without significant death or major impairment of later life. By reversing light cycles, we show that developmental stage differences are not responsible for the circadian muscle growth difference we observe (Fig. 2). Moreover, our data (Fig. 1) indicate that abolition of the circadian clock prevents about a third of the muscle growth that occurs between 1 dpf, when myotomal muscle has just formed, and 3 dpf. Thus, the clock appears to be of importance for muscle growth throughout early larval life.

### A model for circadian- and activity-regulated muscle growth

Our findings suggest a model for the link between muscle protein turnover and muscle growth (Fig. 6). Let us first consider the situation in the absence of activity; growth in both day and night is not detectably altered by rapamycin, which suggests that the regulation of growth is independent of TORC1 signalling (Fig. 4). This view is further supported by the fact that inactive fish show similar pS6/S6 ratio in day and night and have similar RpS6 content (Fig. 4). Nevertheless, the rate of protein synthesis is significantly greater in day than night (Fig. 3), leading to the conclusion that the circadian clock promotes greater protein synthesis during the day compared with the night (Fig. 6, 3^rd^ and 4^th^ columns). On the other hand, in anaesthetised fish, MG132 specifically enhanced the growth at night, but not in the day (Fig. 5), indicating that significant protein catabolism opposing muscle growth only occurs at night. As inactive fish muscle does not grow at night, this implies that the proteasome-dependent night time catabolism has an equal magnitude but opposite effect on muscle growth to nocturnal anabolism (Fig. 6, 4^th^ column). Thus, we hypothesise that the intrinsic growth difference between day and night is due to the presence of rapid anabolism alone in the day (Fig. 6, 3^rd^ column), and the combinatorial presence of slower anabolism and catabolism in the night, where the latter outbalances the former resulting in net zero growth.

**Figure 6.**
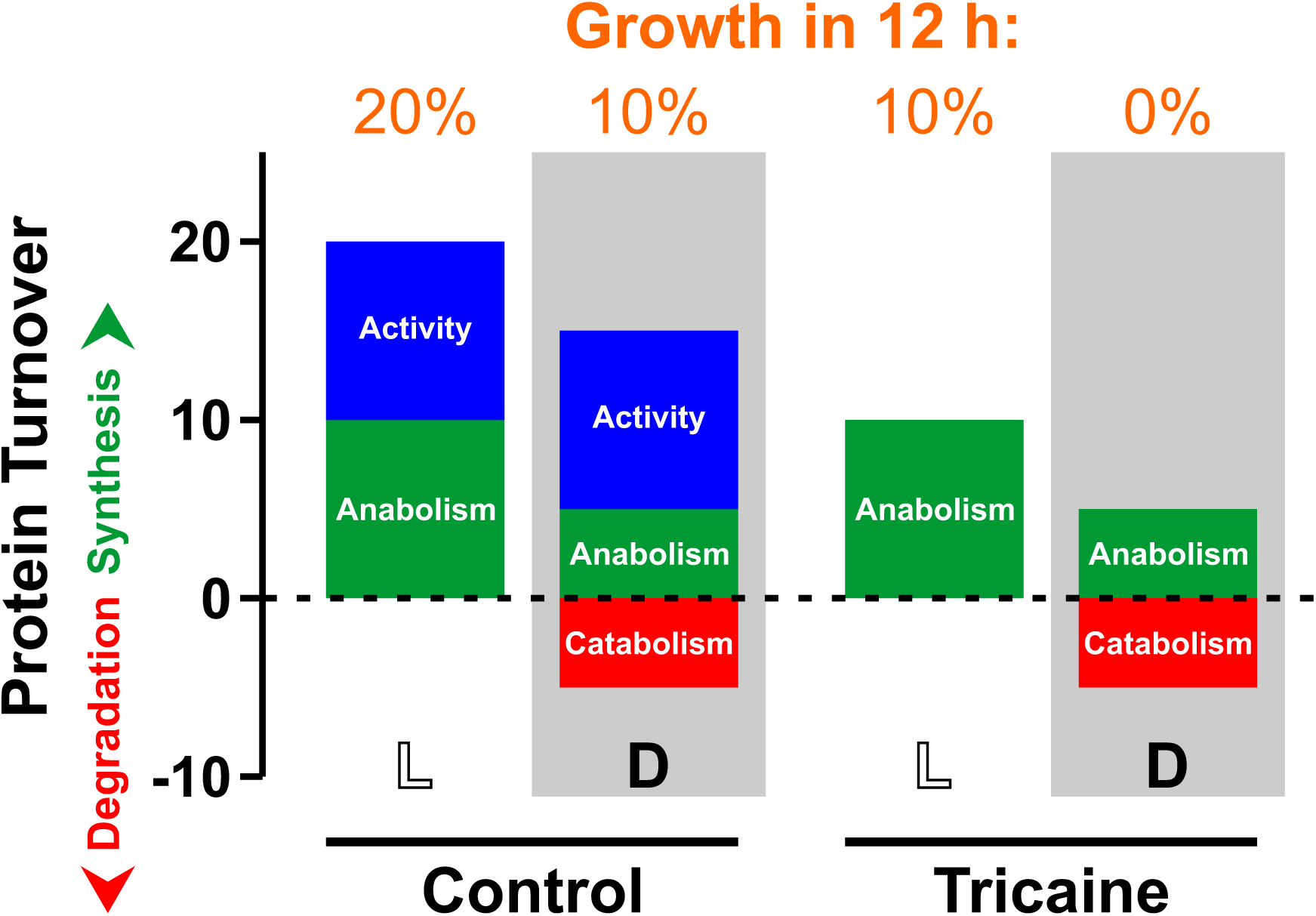
Model of how balance of protein anabolism and catabolism sum to yield observed muscle growth. *Y*-axis represents protein synthesis (positive) and degradation (negative). *X*-axis represents condition: L light/day, D dark/night. Observed (and predicted net) growth over 12 h period is shown above. Blue represents anabolism driven by activity. Green represents activity-independent anabolism that is increased by the circadian clock during the day. Red represents catabolism induced by the circadian clock at night.

Next, consider active muscle at night (Fig. 6, 2^nd^ column); protein synthesis increases significantly compared to inactive muscle at night (Fig. 3), so activity triggers protein synthesis even though pS6/S6 ratio and RpS6 levels appear unaffected (Fig. 4). Congruently, rapamycin has no effect on nocturnal growth (Fig. 4). Yet MG132 still enhances nocturnal muscle growth, just as it did in inactive muscle (Fig. 5), showing that protein catabolism is restricting growth even in active muscle at night.

Lastly, consider active muscle during the day (Fig. 6, 1^st^ column); growth here is fastest, protein synthesis highest and MG132 has no effect (Figs 2,3,5). As the extra growth caused by activity in the day is similar in magnitude to that at night (Fig. 2), and as protein synthesis goes up by a similar amount (Fig. 3), we suggest that the effect of activity is similar to that at night (Fig. 6, 1^st^ and 2^nd^ columns). Yet, strikingly, rapamycin does inhibit growth of active muscle in the day, unlike at night (Fig. 4). However, as the pS6/S6 ratio and total S6 are increased in active muscle during the day (Fig. 4), we propose that the circadian clock alters the response to physical activity by an increase in TORC1 activity that is essential for maximal growth. It remains to be determined if this circadian alteration is intrinsic or extrinsic to the muscle itself.

## Methods

### Zebrafish lines and maintenance

Zebrafish were reared at King’s College London at 28.5 °C with adults kept at 26.5°C on a 14 (09:00-23:00)/10 hours light/dark cycle, with staging and husbandry as described (Westerfield, 2000). Newly-fertilized AB embryos were obtained by pairwise natural spawning at 09:00-10:00 in Tecniplast 1 L mating tanks, cleaned and transferred within 1-2 h to 10 cm triple-vented plastic dishes in incubators at 28.5°C fitted with timed LED lighting systems on a 12 h light (L; 09:00-21:00):12 h dark (D; 21:00-09:00) cycle, or other regimes as described in text. The intensity of light was maintained at around 100 μW/cm^2^ (Tamai et al., 2007). A temperature change of 2°C or more can entrain the circadian clock under otherwise constant conditions (Dekens and Whitmore, 2008). Incubator temperature was monitored with a thermocouple probe every 10 s for 24 h for an entire ligh/dark cycle, revealing a maximum temperature variation of 0.37°C. As the mean temperature in the light/day period was <0.1°C higher than that in the dark/night, any subtle temperature changes are insufficient to entrain the clock under constant light or dark conditions. No more than 60-80 fish were present in each dish and chorion debris, dead or sick fish were removed daily where the light regime permitted. All experiments were performed in accordance with licences held under the UK Animals (Scientific Procedures) Act 1986 and later modifications and conforming to all relevant guidelines and regulations.

### Embryo manipulation

*Tg(Ola.Actb:Hsa.HRAS-EGFP)*^*vu119*^ (Cooper et al., 2005) was maintained by backcrossing on the AB wild-type background. Tricaine (Sigma Aldrich) was prepared as described (Westerfield, 2000). Rapamycin (Enzo Life Sciences) and MG132 (AlfaAesar) were prepared in DMSO at stock concentrations of 5 mM and 1 mM, respectively, stored at −20°C and diluted to 10 μM in either fish water alone or tricaine-containing fish water when used. Control groups were treated with 0.2-1% DMSO alone.

### O-propargyl-puromycin (OPP) protein synthesis assay

Experiments were performed using Click-iT^®^ Plus OPP Protein Synthesis Assay Kit with Alexa Fluor^®^ 488 picolyl azide (Invitrogen). Briefly, larvae were treated with 50 μM Click-iT^®^ OPP Reagent (Invitrogen) for 2 h at either 3 dpf (ZT0-12) or 3.5 dpf (ZT12-14), before fixing with 2% paraformaldehyde for 30 min at room temperature. As a negative control, larvae were treated with 200 μg/mL cycloheximide (Sigma Aldrich) for 1 h prior to labelling. Fixed larvae were bleached in 10% H_2_O_2_ and 5% formamide solution to remove pigment as described (Diogo et al., 2008), and then permeabilised with phosphate-buffered saline (PBS) containing 0.1% Triton X-100 and 1% DMSO. Permeabilised larvae were then labelled with Click-iT^®^ Plus OPP reaction cocktail (Invitrogen) for 1 h in dark in room temperature according to manufacturer’s instructions. Labelled larvae were then washed thoroughly with PBS, mounted and subjected to confocal imaging with no saturated or black pixels. Within an experiment, all fish were imaged on identical settings. Quantification of the mean fluorescent intensity was performed on single optical sections at equivalent depth in whole somite 17 using Fiji (NIH). Normalised signals were calculated by subtracting mean signals measured in negative control (cycloheximide) samples at ZT0-2.

### Confocal imaging

For confocal imaging, larvae were mounted in 0.8% low melting point agarose, and data collected on the somites 17-18 near the anal vent on a LSM 5 Exciter microscope (Zeiss) equipped with 20x/1.0W objective and subsequently processed using either Volocity 6.3 (Perkin Elmer), Fiji (NIH) or ZEN (Zeiss) software. For live imaging, mounted larvae were transiently anaesthetised with tricaine. Immediately after the scan, larvae were washed and then separately housed in 24-well or 96-well plates to track individual myotome growth. Medium in each well was changed every 12 h. Myotome volume was calculated as described (Hinits et al., 2011; Roy et al., 2017) and schematised in Fig. S1A.

### Protein extraction, SDS-PAGE, and Western blotting

Total protein was extracted from pools of twenty larvae via sonication in solution mixtures containing Tissue Extraction Reagent I (Invitrogen), cOmplete™, EDTA-free Protease Inhibitor Cocktail Tablets (Sigma Aldrich), and phenylmethylsulfonyl fluoride (PMSF). Protein extracts were then pelleted by centrifugation, after which the supernatant was mixed with Laemmli Sample Buffer (Bio-rad) and 2-mercaptoethanol (Bio-rad) and heated at 95°C for 5-10 min, before subjecting to SDS-PAGE analysis. Protein extract equivalent to 1-2 larvae was loaded per lane. SDS-PAGE was performed using Mini-PROTEAN^®^ TGX Precast Gels (Bio-rad), and separated proteins were then transferred to nitrocellulose membranes, blocked in either 5% non-fat dry milk or 5% BSA in Tris-buffered saline (TBST), incubated in primary and secondary antibodies at 4°C overnight and at room temperature for 1 h, respectively. Primary antibodies used were: S6 ribosomal protein (1:1000; #2317; Cell Signaling), phospho-S6 ribosomal protein (1:1000; #5364; Cell Signaling), and histone H3 protein (1:1000; #ab1791; Abcam). Secondary antibodies used were: HRP GoatαMouse IgG(H+L) (1:2000; #AP308P; Sigma Aldrich) and Alexa Fluor 488 GoatαRabbit IgG(H+L) (1:2000; #A28175; Invitrogen). Signal detection was performed using ChemiDoc™ Imaging System (Bio-rad) and analysed on Image Lab™ Software (Bio-rad). pS6/S6 ratio was normalised to control samples at ZT3.

### RNA extraction, RT-PCR and qPCR

Total RNA was extracted from pools of five larvae using Tri-reagent (Sigma Aldrich) via sonication, isolated via phase separation using 1-bromo-3-chloropropane, precipitated using 2-propanol and washed using 75% ethanol. Extracted RNA was treated with TURBO DNase (Invitrogen) for 15-30 min at 37°C. Purified total RNA (500 ng) was then reverse transcribed using Eurogentec Reverse Transcriptase Core Kit (Eurogentec) with both random and oligo-dT primers following manufacturer’s instructions. qPCR was performed and results analysed as described (Ganassi et al., 2018). Briefly, technical triplicates were performed on 5 ng of cDNA using Takyon Low ROX SYBR 2X MasterMix blue dTTP (Eurogentec) on a ViiA™7 thermal cycler (Applied Biosystems). ΔCt was calculated by subtracting the Ct value of the *ef1a* housekeeping gene from that of the target gene. ΔΔCt of each target gene was then calculated by subtracting the average ΔCt of samples, and relative gene expression calculated as 2^-ΔΔCt^ (Livak and Schmittgen, 2001), followed by normalising to control samples at ZT3. Primers used are listed in Table S1.

## Statistical analyses

Quantitative analysis on images was performed using Fiji (NIH). Statistical analysis was performed on Prism 8.2.1 (GraphPad). The test for normality was performed on myotome volume measured at 3 dpf and 4 dpf using D’Agostino and Pearson test (Fig. S2C,D). *p*-values >0.05 indicate that the data are not inconsistent with a Gaussian (normal) distribution. We, therefore, presume that all further myotome volume measurements are normally distributed, justifying use of ANOVA with two factors (‘experiment/day’ and ‘treatment’) followed by Bonferroni’s post-hoc test to assess differences. All data were shown with bars representing mean ± standard error of the mean (SEM). All experiments were performed on at least three independent biological replicates. *p*-values for rejection of the null hypothesis of no difference between groups are indicated above lines indicating the columns compared, and are given in the form XE-Y, meaning X × 10^−Y^. For the detection of rhythmic components in qPCR data sets, JTK_Cycle, a non-parametric algorithm (Hughes et al., 2010) was applied, using the statistical computing software R version 3.5.2.

## Data availability

The authors declare that all data supporting the findings of this study are available within the article and its supplementary information files or from the authors upon request.

## Author contribution

JJK and SMH conceived the project and designed experiments. TGP performed experiments and corresponding analysis depicted in Figures 1A and S2A,B. JJK performed all the other experiments and analysis. SMH and JJK wrote the paper. SMH obtained finance.

## Acknowledgements

We thank our Randall colleagues, particularly Massimo Ganassi for advice, Fiona Wardle for anti-H3 antibody and the KCL fish facility staff for their support. SMH is a Medical Research Council Scientist with MRC Programme Grant MR/N021231/1 support.

---

**Supplementary Figure S1.**
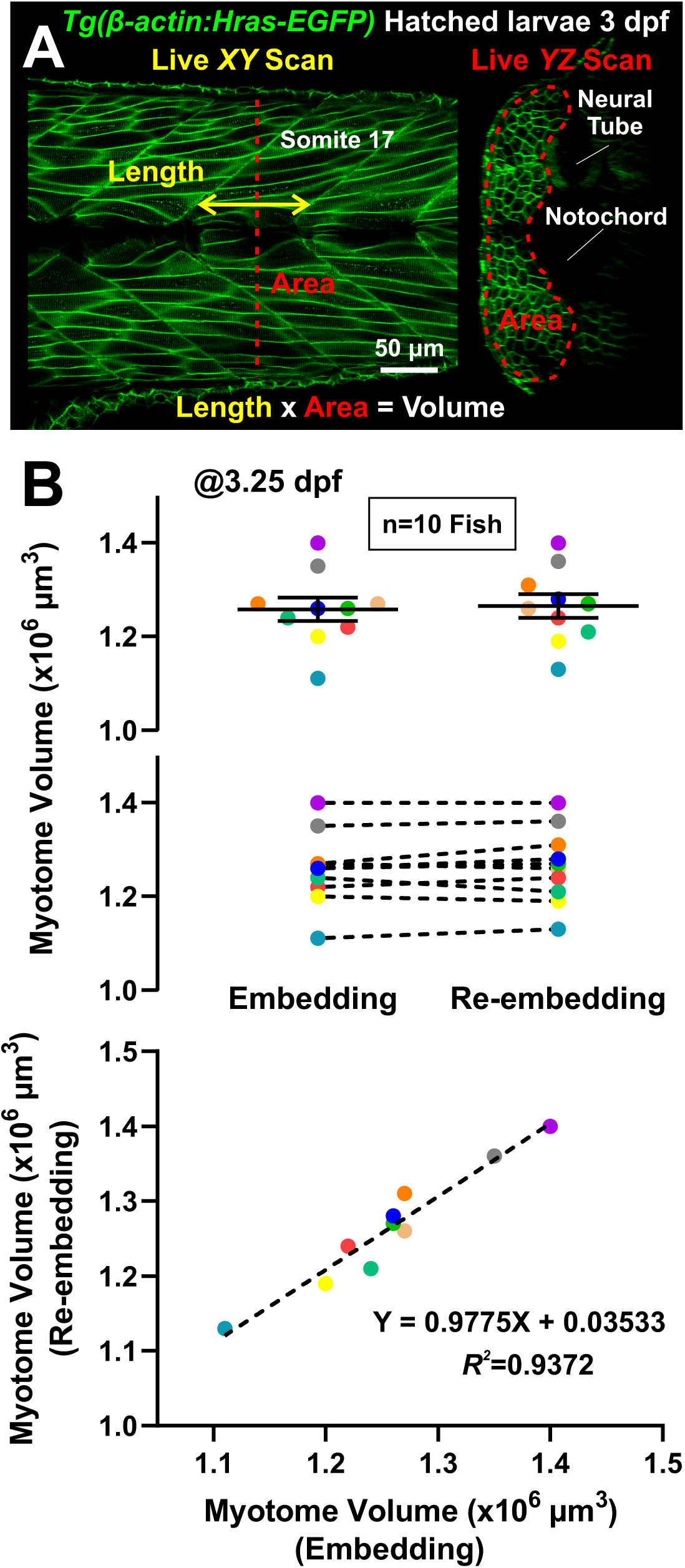
Live muscle volume assay is highly reproducible. **A.** *β-actin:Hras-EGFP* transgenic larvae were transiently anaesthetised, mounted, and scanned from left lateral view (dorsal to top) in *XY* and *YZ* optical confocal sections of somite 17 to measure the size of the myotome. **B.** Repeated re-embedding and re-scanning of single larvae within 1 h (individuals indicated by colour) showed high concordance (<3% variation, R=0.94) in measured myotome volume (n=10 fish). Note the slight trend towards increase in mean mass, consistent with growth between the two daytime scans.

**Supplementary Figure S2.**
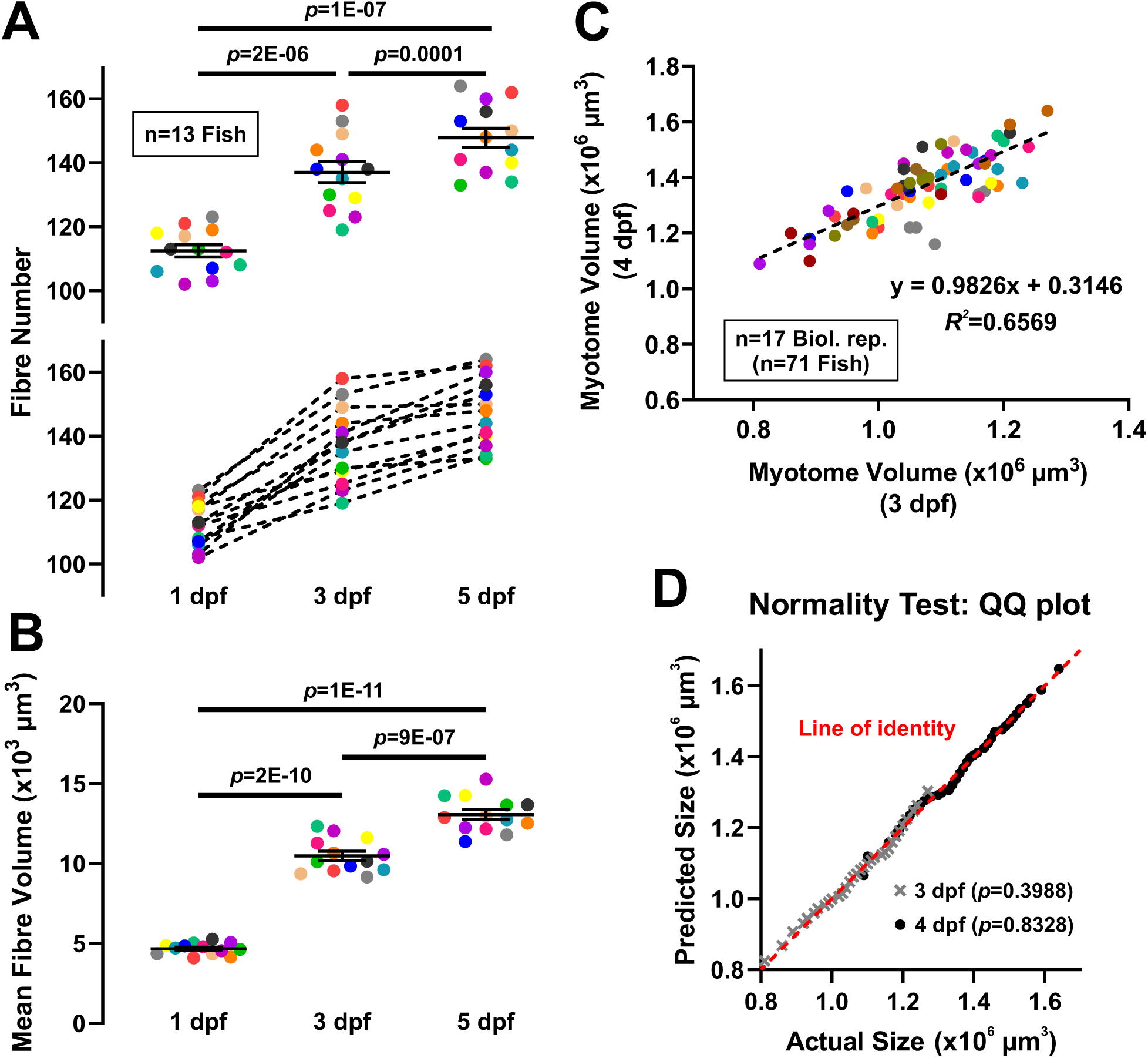
Muscle grows by increase in both fibre number and fibre volume. Individual embryos/larvae were repeatedly imaged and scanned to assess variation in growth rates between individuals and lays. Somite 17 myotome volume and fibre number were determined. **A.** Two representations of the fibre numbers in the same individual fish (indicated by colour, corresponding to the myotome volume data shown in Fig. 1B) showing the increase from 1-5 dpf (n=13 fish)**. B.** Increase in mean fibre volume (calculated as [myotome volume]/[fibre number]) as individual fish grow (same colour code as Fig. 1B and Fig. S1A). **C.** Myotome volume of individual larvae at 3 dpf were compared with the same individuals at 4 dpf (n=71 fish, from 17 biological replicates). Colours represent different lays. **D.** QQ plot showing lack of deviation from a Gaussian (normal) distribution as analysed by the D’Agostino and Pearson normality test, where the null hypothesis is that all the values were sampled from a population that follows a Gaussian distribution. *p*-values >0.05 indicate the data are not inconsistent with a Gaussian distribution. Symbols distinguish myotome volume measure at 3 dpf and 4 dpf.

**Supplementary Figure S3.**
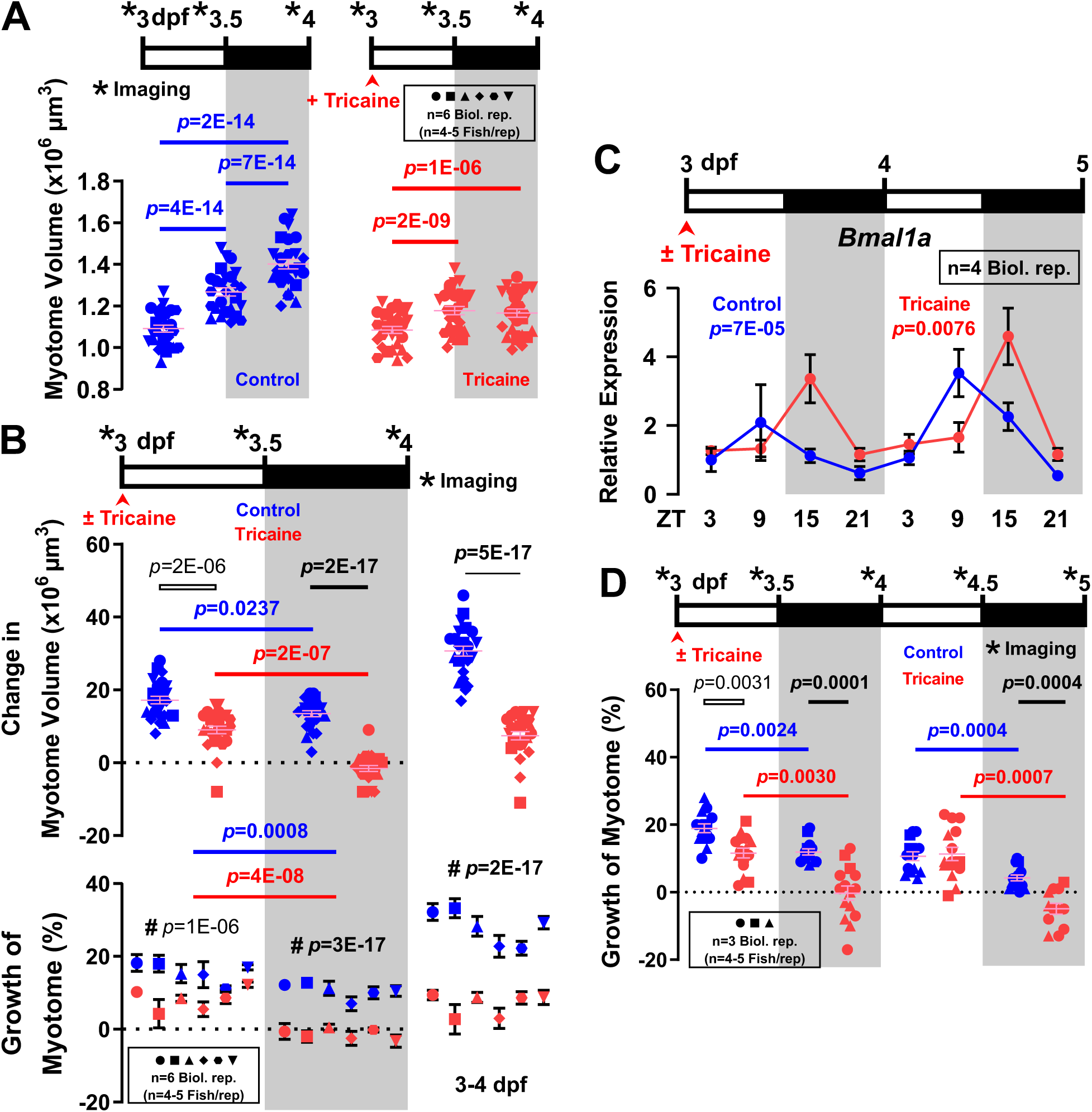
Circadian growth difference in absolute myotome volume across lays and developmental stages. **A**,**B.** Data underlying that presented in Fig. 2A as myotome growth (%). Larvae raised under LD (n=25-26 fish from 6 biological replicates) were either untreated (Control, blue) or anaesthetised (Tricaine, red) from 3-4 dpf. Confocal imaging was performed every 12 h (*) over the 24 h period to measure myotome volume. Symbols shapes distinguish biological replicates from separate lays. Colours represent different treatments. Graphs show myotome volume of individual fish at each timepoint (A), absolute change in myotome volume for each fish across all six lays (B upper) and mean±SEM for each individual lay (B lower). Note the consistency in growth and effects of time and anaesthetic, despite differences in absolute mean size between lays. **C.** Expression of core clock gene *bmal1a* was assayed by qPCR on total RNA collected every 6 h from untreated (Control, blue) and 3-5 dpf anaesthetised (Tricaine, red) larvae raised under LD (n=4 biological replicates). *Ef1a* was used for normalisation. Statistic represents JTK_Cycle (see Methods). **D.** Growth of myotome (%) of untreated (Control, blue) and 3-5 dpf anaesthetised (Tricaine, red) larvae raised under LD (n=12-15 fish from 3 biological replicates). Note the declining growth rate with age, consistent effect of anaesthetic and persistent circadian difference with or without anaesthetic.

**Supplementary Figure S4.**
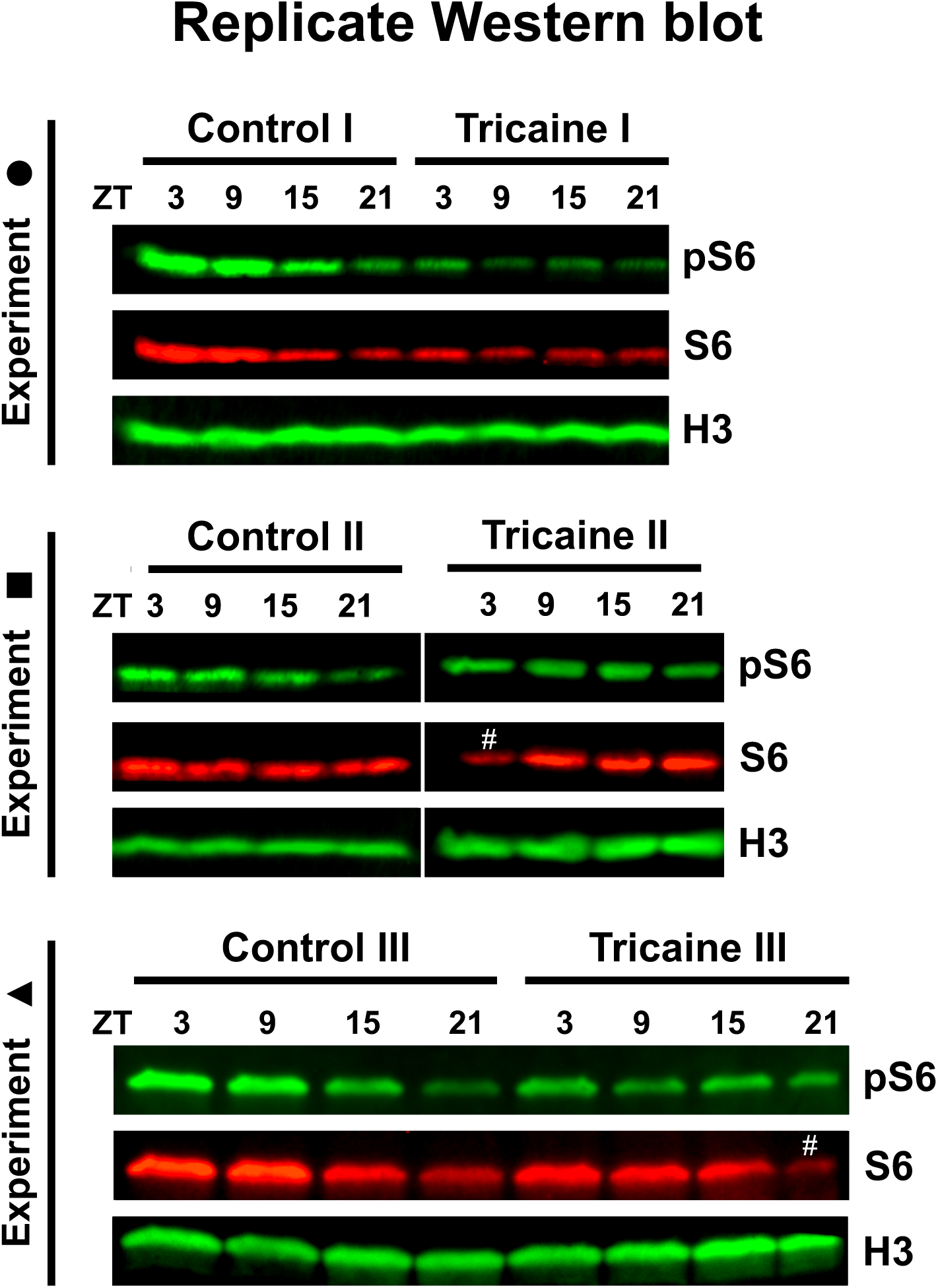
Circadian rhythm and activity regulate RpS6 phosphorylation. Detection of phosphor-Ribosomal protein S6 (pS6), total RpS6 (S6) and Histone H3 loading control (H3) in replicate Western blots (including that shown in Fig. 4A) from three separate lays (I-III; shape symbols correspond to data presented in Fig. 4B) of fish grown in fish water (Control) or fish water plus Tricaine, consistently showing a) increased S6 at ZT3 in control, b) decreasing pS6/S6 ratio in control at ZT21 and c) ablation of daytime pS6/S6 increase by tricaine. Note that the two highest points in tricaine (indicated by # and quantified in Fig. 4B) have unusually weak S6 bands rather than high pS6 bands, possibly indicating an S6 detection problem. Data in Experiment II are from two gels run and blotted in parallel.

**Table S1.**
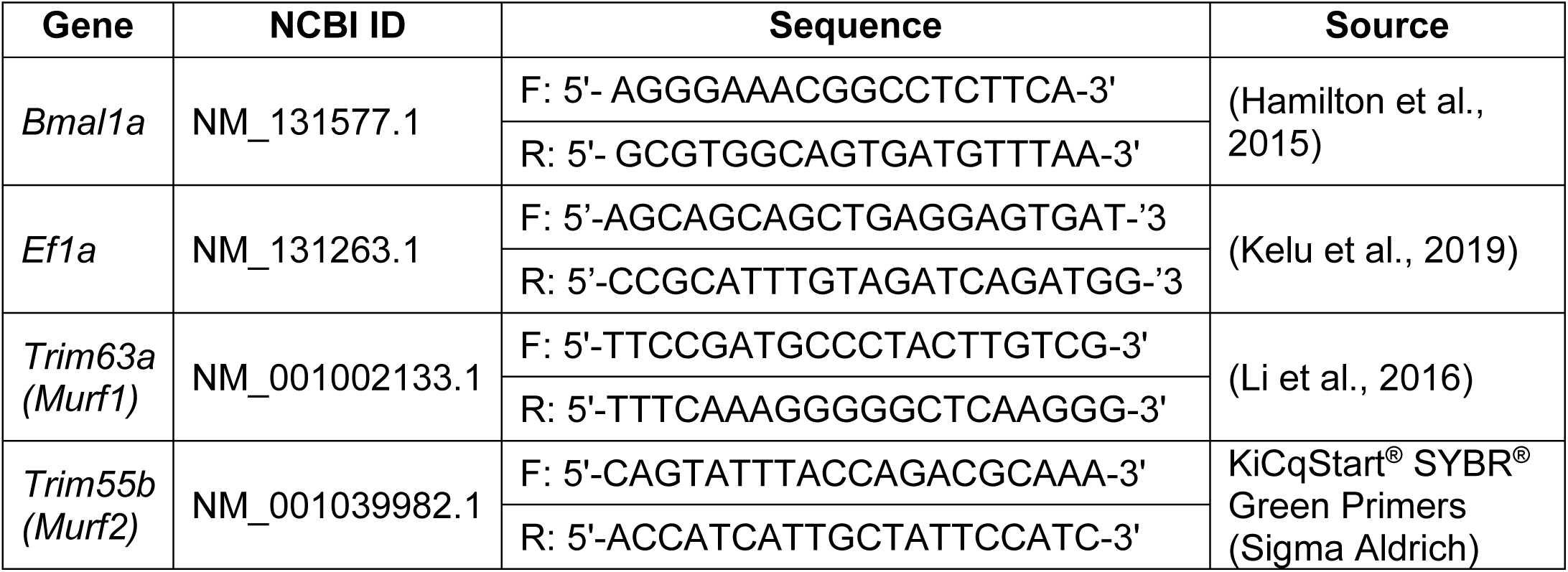
Primer used.

